# Perpendicular axes of incipient speciation generated by mitochondrial introgression

**DOI:** 10.1101/072942

**Authors:** Hernán E. Morales, Paul Sunnucks, Leo Joseph, Alexandra Pavlova

**Affiliations:** School of Biological Sciences Monash University, Clayton Campus, Melbourne, 3800 Victoria, Australia; Department of Marine Science, University of Gothenburg, Medicinaregatan 9C, 41390 Gothenburg, Sweden; Australian National Wildlife Collection, CSIRO National Research Collections Australia, GPO Box 1700, Canberra ACT 2601

**Keywords:** mitochondria, mitonuclear, selection, introgression, speciation, selective sweep

## Abstract

Differential introgression of mitochondrial versus nuclear DNA generates discordant patterns of geographic variation and can promote speciation. We examined a potential case of mitochondrial introgression leading to two perpendicular axes of differentiation. The Eastern Yellow Robin, a widespread Australian bird, shows a deep mitochondrial split that is perpendicular to north-south nuclear DNA and plumage colour differentiation. We proposed a scenario to explain this pattern: (1) the two nuclear and mitochondrial genomes differentiated in concert during north-south population divergence; (2) later, their histories disconnected after two mitochondrial introgression events resulting in a deep mitochondrial split perpendicular to the nuclear DNA structure. We tested this scenario by coalescent modelling of ten mitochondrial genes and 400 nuclear DNA loci. Initial mitochondrial and nuclear genome divergences were estimated to have occurred in the early Pleistocene, consistent with the proposed scenario. Subsequent climatic transitions may have driven later mitochondrial introgression. We reject neutral introgression and consider evidence consistent with adaptive mitochondrial introgression and selection against incompatible mitochondrial-nuclear combinations. This likely generated an axis of incipient speciation associated with mitochondrial differentiation in the face of nuclear gene flow, perpendicular to the initial north-south axis of incipient speciation (reflected in nuclear differentiation and colour variation).

## Introduction

When divergent populations undergo hybridization, genes from one population can be incorporated into the other (i.e., there is introgression) to a variable extent across the genome (Harrison & Larson, 2014; Harrison & Larson, 2016). Alleles that are not involved in local adaptation and that have not accumulated incompatibilities with other loci are expected to move freely between populations (Mallet, 2005). On the other hand, if genes from one population improve fitness in the other population, adaptive introgression can occur (Hedrick, 2013). The proportion of the genome that is resistant or prone to introgression can vary as a result of local adaptation in heterogeneous environments and demographic history (Harrison & Larson, 2014). Therefore, differential rates of introgression offer a valuable insight into adaptive divergence and speciation (Payseur, 2010; Rheindt & Edwards, 2011; Toews & Brelsford, 2012).

Differential rates of introgression of mitochondrial DNA (mtDNA) versus nuclear DNA (nDNA) genes are a main cause of mitochondrial-nuclear (mitonuclear) discordances (Toews & Brelsford, 2012). However, it is challenging to predict the conditions under which higher rates of mitochondrial or nuclear introgression can be expected. This is because genetic patterns in both genomes can differently reflect the effects of genetic drift and selection, and the two genomes have different modes of inheritance and recombination (Harrison, 1990; Funk & Omland, 2003). Under selective neutrality, the rate of allelic replacement should be lower for genetic markers having higher intraspecific gene flow and associated with the more dispersive sex (Currat *et al.*, 2008). Thus, under female-biased dispersal, mtDNA introgression should be lower than that of nDNA, an idea supported by meta-analysis (Petit & Excoffier, 2009). Congruently, low passage through hybrid zones of maternally-transmitted mtDNA is predicted for species with heterogametic females, such as birds, under Haldane’s Rule (i.e. disproportional hybrid sterility and/or inviability of the heterogametic sex; Haldane, 1922). These predictions are commonly supported by studies of avian hybrid zones (Rheindt & Edwards, 2011). However, birds do not always have lower mtDNA than nDNA introgression. Potential reasons for higher mtDNA than nDNA introgression, other than male-biased dispersal (rare in birds), include strong demographic disparities between divergent populations, and adaptive introgression (Currat *et al.*, 2008; Toews & Brelsford, 2012; Hedrick, 2013). Given the importance of mtDNA for organismal metabolism and fitness, adaptive mitochondrial introgression into a beneficiary population could be common (Dowling *et al.*, 2008). Thus, bird species with female-biased dispersal that show greater mitochondrial than nuclear DNA introgression are strong candidates for adaptive introgression.

The Eastern Yellow Robin (*Eopsaltria australis*, hereafter EYR) shows a striking pattern of geographic mitochondrial-nuclear (mitonuclear) discordance, representing an excellent system to study differential mtDNA and nDNA introgression (Pavlova *et al.*, 2013). The two major mitochondrial lineages of EYR (mitolineages; mito-A and mito-B) are 6.8% divergent and structured across inland and coastal sides of the Great Dividing Range in south-eastern Australia (Pavlova *et al.*, 2013; Fig. 1A). In contrast, the major axis of nDNA structure runs north-south through the species range (Fig. 1B). Thus, nDNA and mtDNA structures are geographically perpendicular (Pavlova *et al.*, 2013; Fig. 1A-B). Additional, minor inland-coastal nDNA structure exists in the south corresponding with mitolineage distributions (Morales et al., submitted; Fig. 1B). The major north-south axis of nDNA differentiation is mirrored by rump plumage colour variation, supporting two currently recognized subspecies: the rump is bright yellow in northern *E. a. chrysorrhoa*, and olive-green in southern *E. a. australis* (Ford, 1979; Schodde & Mason, 1999). Colour variation at the continental-scale is strongly influenced by population history, but on a regional scale is structured according to local environmental variation (Morales *et al*., submitted).

**Figure 1.**
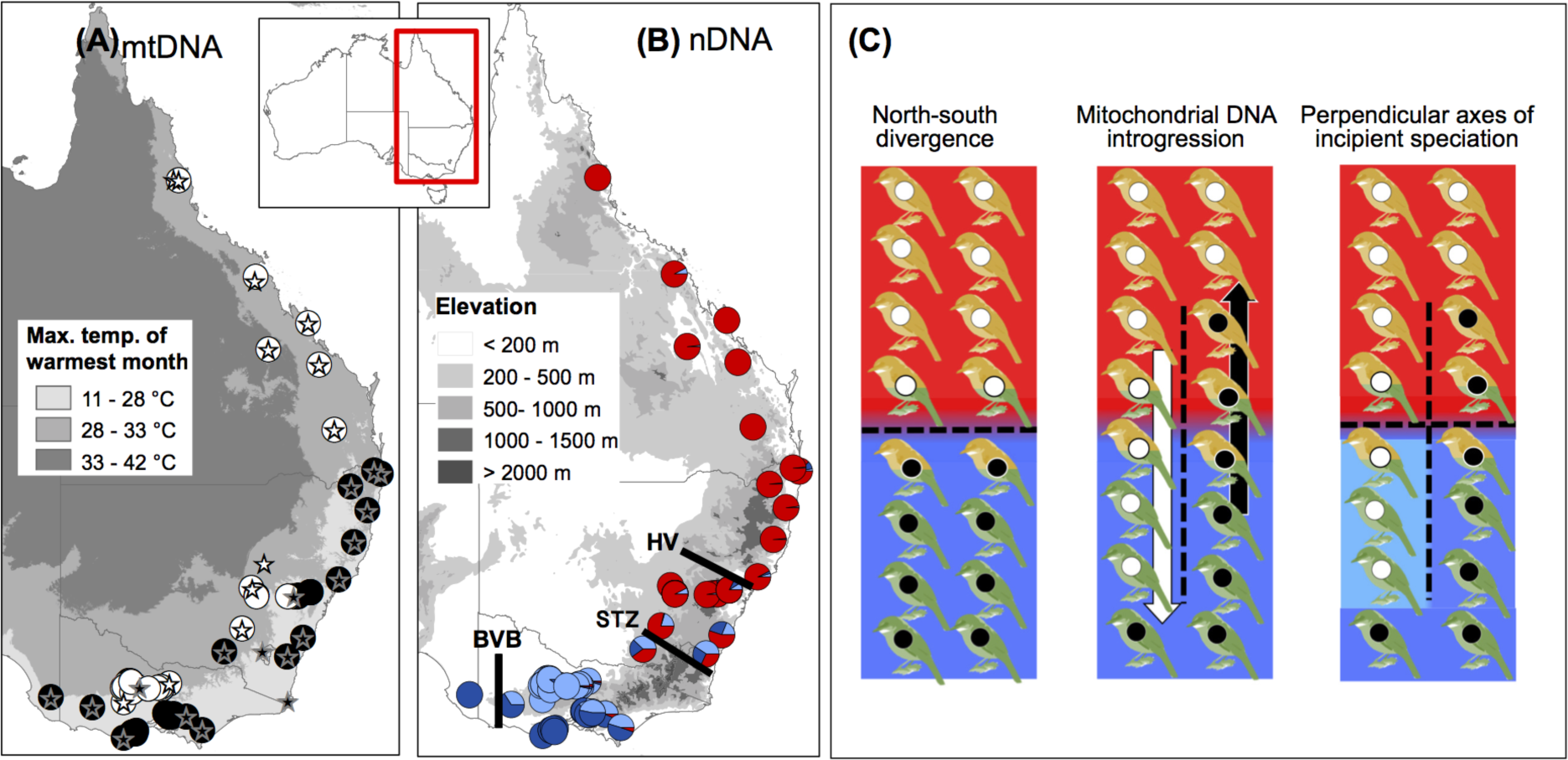
Distribution of Eastern Yellow Robin samples used in this study showing their contribution to mitochondrial (A) and nuclear (B) genetic structures and a schematic representation of EYR evolutionary history (C). (A). Distribution of mitochondrial lineages mito-A (white circles) and mito-B (black) plotted over the maximum temperature of the warmest month. Circles represent samples sequenced for the mitochondrial ND2 gene, stars-samples for which data from 10 mitochondrial genes were used. The northernmost sample of mito-A mitolineage displays late-Pleistocene intralineage divergence (Fig. S7). (B) Pies-membership in three genetic clusters according to *K*=3 STRUCTURE analysis (see Fig. 2A): northern cluster (red), southern coastal cluster (dark blue) and southern inland cluster (light blue). The background represents an elevation map featuring the Great Dividing Range dividing the mitochondrial lineages (see panel B). Black lines represent potential vicariant/environmental barriers, HV-Hunter Valley, STZ-Southern Transition Zone, and BVB-Bassian Volcanic Barrier. (C) Evolutionary history of the Eastern Yellow Robin, the colour of the boxes represent their nuclear genomic background (colour of the background; north = red and south = blue), their plumage colouration (colour of the birds; northern yellow, southern green and intermediates of mixed colour), and their mitochondrial membership (colour of the circles; mito-A = white and mito-B = black). First panel shows the first axis of incipient speciation: mtDNA, nDNA and colour differentiation between northern and southern birds with a zone of intergradation. The second panel shows the second axis of incipient speciation: two independent events of mitochondrial introgression occurred without nDNA introgression, resulting in mtDNA genetic structure in the inland-coastal direction. The third panel shows the current perpendicular pattern of differentiation: inland-coast mitochondrial divergence in the face of nuclear gene flow, major north-south nDNA structure and plumage colouration divergence and a minor inland-coast nuclear DNA divergence in the southern range (shades of blue).

Previous studies considered drivers of observed patterns of genetic and phenotypic variation in EYR. Using microsatellite, nuclear intron data and one mitochondrial gene, Pavlova *et al.* (2013) rejected three common explanations of mitonuclear discordance based on selective neutrality (Toews & Brelsford, 2012): (1) inland-coast vicariance was not supported by models of past and present species distribution, (2) incomplete lineage sorting was contradicted by the >1500 km extent of the mitochondrial split and inferred nuclear gene flow between mitolineages, and (3) male-biased dispersal is counter to known female-biased dispersal in EYR (Debus & Ford, 2012). They found that maximum temperature of the hottest month explains mtDNA variance over and above that explained by geographic position and distance, which suggests environmental temperature as a selective driver of mitolineage distribution (Fig. 1A). Pavlova *et al.* (2013) concluded that the major nDNA north-south structure in EYR was consistent with isolation-by-distance, and that inland-coastal mtDNA divergence occurred *in situ*. Subsequently, Morales *et al.*, (2015) found supporting evidence for selection on mitochondrial genomes and confirmed extremely low mitogenome-wide intra-lineage diversity consistent with selective sweeps. Morales et al. (submitted; Fig. 1B) expanded the analysis of nDNA by analysing genome-wide neutral single nucleotide polymorphisms (SNPs) and found two northern and southern genetic clusters with a narrow zone of intergradation, contrary to isolation-by-distance. They also showed that plumage colour differentiation follows a similar geographic trend, albeit with a broader zone of intergradation. Reconstructions of the evolutionary histories of each genome are needed to better understand mitonuclear discordance in EYR.

Here we propose a novel scenario to explain perpendicular mitonuclear differentiation in EYR (Fig. 1C). Initial north-south divergence generated concerted mtDNA and nDNA divergence, currently reflected in the major nDNA structure and colour variation (first axis of incipient speciation) but not in mtDNA structure. Subsequently independent events of mitochondrial introgression might have occurred without nDNA introgression, one south-to-north coastwards of the Great Dividing Range, and the other north-to-south inland of the Great Dividing Range. Mitochondrial introgression would then have resulted in the current inland-coastal mitochondrial split and inland-coast mitonuclear divergence-with-gene-flow in the southern population (a second axis of incipient speciation). We used a coalescent multilocus approach to test this scenario by analysing 10 mitochondrial genes free from signatures of selection, and 400 sequenced nuclear loci (Peters *et al.*, 2007; Rheindt & Edwards, 2011). We estimated nuclear divergence times, gene flow rates and effective population sizes and tested whether the onset of mitochondrial divergence coincided with north-south population divergence. We discuss how adaptive mitochondrial evolution, introgression, and mitonuclear co-evolution could have driven EYR divergence and led to incipient speciation (Dowling *et al.*, 2008; Gershoni *et al.*, 2009; Burton *et al.*, 2013; Hill, 2015; Hill, 2016).

## Methods

### Samples, molecular methods and datasets

We analyzed (1) mitochondrial ND2 sequences, (2) 2728 SNPs and (3) phased alleles for 400 nuclear sequences for 69 individuals, and (4) 10 mitochondrial genes for 32 individuals (Fig. 1A-1B). Genomic DNA from 42 newly collected blood samples was extracted with DNAeasy Kit (Qiagen, Germany) following the manufacturer’s protocol. For these samples, a partial region (∼1000 bp) of mitochondrial ND2 gene was amplified following Pavlova *et al.* (2013) and sequenced commercially (Macrogen, Korea). The newly produced ND2 dataset was supplemented with previously published ND2 sequences for 27 individuals (Genebank accession in Table S1; Pavlova *et al*., 2013). Based on ND2, all 69 individuals were assigned to one of the two mitolineages (35 mito-A and 34 mito-B; Table S1; black and white circles on Fig. 1A).

For the same 69 individuals, 1000 sequenced anonymous nuclear loci were obtained by hybrid capture enrichment probes (size = 240 bp; Lemmon *et al.*, 2012; Morales *et al*., submitted) and analyzed as a set of 2728 SNPs, previously used by Morales *et al*., (submitted), and as phased alleles. Details of the hybrid capture enrichment method are in Appendix S1. More details about post-processing of targeted captured loci including raw sequencing reads processing, read assembly, orthology calculations and sequence alignment are in Prum *et al.* (2015) with accompanying generic scripts on doi:10.5281/zenodo.28343. Allele phasing for each locus was determined statistically from the assembled reads by drawing a posterior distribution for each individual separately, following Pyron *et al.* (2016); for methodological details and scripts see doi:10.5061/ dryad.51v22. In short, the method generates alleles with no ambiguities for those positions that can be phased with a ≥ 95% posterior probability confidence, leaving ambiguities for the remainder of the polymorphic sites.

To appropriately assign inheritance scalars for the sequenced loci in the IMa2 analysis (below), we mapped the alignment consensus sequences to the Zebra Finch *Taeniopygia guttata* genome taeGut3.2.4 (Warren *et al.*, 2010) using BLASTn v.2.3.0 (Camacho *et al.*, 2009) with a E-value threshold of 1 × 10^−4^ (BLAST output doi: 10.6084/m9.figshare.3581004). Because historical inferences assume neutral evolution of markers, we identified all nuclear loci that may have evolved under directional selection and removed them from SNP and sequenced markers datasets (Appendix S1). From the latter dataset, 400 loci (the maximum number accepted by IMa2, below) were randomly selected for coalescent analyses. Data used in this project (capture probes for the EYR hybrid capture enrichment, raw reads, and phased sequence alignments) have been deposited in figshare (doi:10.6084/m9.figshare.3581004).

Sequences of 10 protein-coding mitochondrial genes (ND1, ND2, ND3, ND5, ND6, COX1, COX2, COX3, ATP6 and ATP8) were extracted from 32 published mitogenome sequences (14 mito-A and 18 mito-B, represented by stars on Fig. 1A; Genbank accessions in Table S2; Morales *et al*., 2015). These genes were chosen because they did not show signatures of positive selection between mitolineages (Morales *et al.*, 2015).

### Nuclear DNA genetic structure

We used the SNP dataset and admixture model with correlated allele frequencies implemented in STRUCTURE 2.3.4 (Pritchard *et al.*, 2000) to confirm the presence of major and minor nDNA structure and assign each individual to one of three populations (northern, south-inland and south-coast: red, light blue and dark blue on Fig. 1C, respectively) for IMa2 analysis (below). STRUCTURE analysis was performed on 706 unlinked SNPs, assuming 1 to 5 genetic clusters (*K*) with 25 Markov chain replicates of 200,000 iterations of burn-in and 80,000 recorded iterations for each *K* (see Supplementary Methods for details). Individuals with a posterior probability (Q-values) of ≥0.8 of belonging to a particular cluster were assigned to that genetic cluster (i.e. population).

We used discriminant analysis of principal components (DAPC: Jombart *et al.*, 2010) implemented in adegenet 2.0.0 for R 3.1.2 (Jombart & Ahmed, 2011; R Development Core Team, 2014) to estimate the amount of genetic variation explained by the major and minor axes of nDNA structure. The DAPC analysis was conducted with 10 populations delimited based on their geographic location (Fig. 2B).

**Figure 2.**
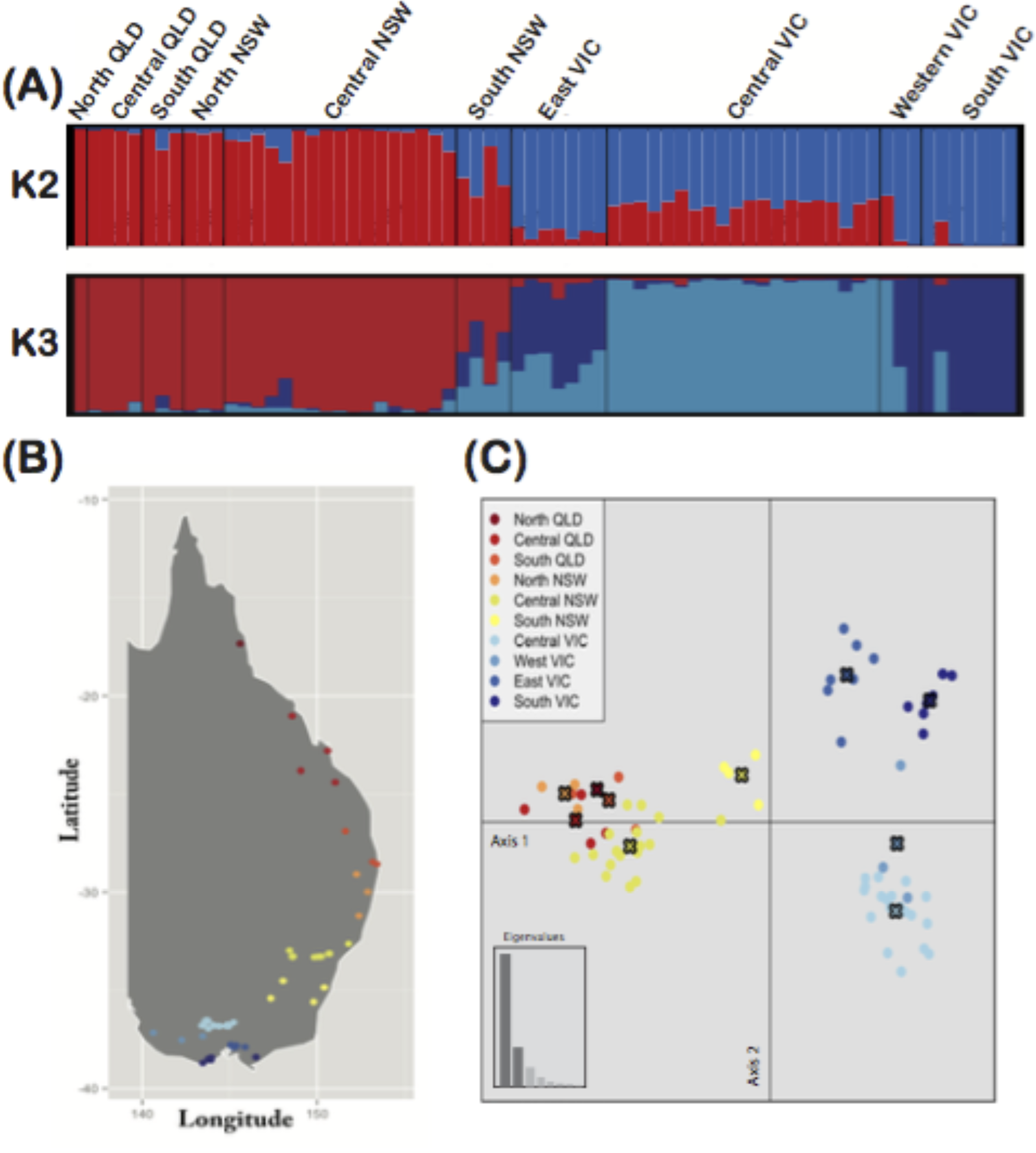
Summary of population genetic structure in the Eastern Yellow Robin. (A) The results of STRUCTURE (Pritchard *et al.*, 2000) models when *K* = 2 and *K*=3. Results were summarised with STRUCTURE HARVESTER Web v0.6.94 (Earl, 2012) and CLUMPP v 1.1.2 (Jakobsson & Rosenberg, 2007). (B) Map of samples used for the nuclear DNA data arranged into arbitrary populations according to geographic position. (C) Discriminant Analysis of Principal Components (DAPC; Jombart *et al.*, 2010) for populations from panel B, axis 1 (PC1) captured 48% of genetic variation and axis 2 (PC2) captured 12% of genetic variation.

### Isolation-with-migration models

To estimate times of population divergence, gene flow and effective population sizes, we fitted a three-population model of isolation-with-migration (Hey & Nielsen, 2004) implemented in IMa2 (Hey, 2010) to the multilocus dataset comprising sequences of 400 randomly-selected anonymous nuclear loci. For each locus, the longest stretch of each sequence without ambiguities was used. Recombination points were detected for each locus with the program IMgc (Woerner *et al.*, 2007) and the longest non-recombining block was retained. Resulting alignments had lengths from 124 to 529 bp (mean = 264 bp). IMa2 estimates model parameters scaled by mutation rate (µ): population divergence time in generations (*t* = tµ), migration rate (*m*=m/µ) and effective population sizes (θ=4N_e_µ).

For efficient computation in the IMa2 analysis, we subsampled individuals from each population (based on Q>0.8 from K=3 Structure analysis; northern: N=18 alleles, southern inland: N=14 and southern coastal: N=16, Table S1). Individuals with a posterior probability (Q-values) of cluster membership <0.6 were not used in the analysis, se we might have underestimated gene flow, however, precise estimates of gene flow are not crucial given our aims. Sixteen parameters were estimated: two of *t*, eight of *m* and six of θ (θs were estimated at three time intervals: *t*1, *t*0 and present; model option j5). Multiple preliminary analyses were run to optimize prior parameter boundaries. Eight independent replicates were run for the final analysis, each involved > 11 × 10^6^ steps after a burn-in period of > 4.8 × 10^5^ steps, employing 150 MCMC chains with geometric heating (parameters a = 0.999 and b = 0.3). Parameter estimates were converted to demographic units using a generation time of 3.5 years (Pavlova *et al*., 2013) and a mutation rate of 1.2 x10^−9^ (lower bound = 0.7 × 10^−9^; upper bound = 2.5 × 10^−9^) nucleotides per base per year (Ellegren, 2007; Lee & Edwards, 2008).

### Mitochondrial lineage divergence

To test for simultaneous divergence of mitolineages (mito-A and mito-B) and nuclear DNA, we built a calibrated phylogeny in BEAST v.1.8.0 (Drummond *et al.*, 2012) using sequences of 10 protein-coding mitochondrial genes that are free from signatures of positive selection. Although mitolineage divergence time was estimated previously from ND2 (1.5 (0.98–2.15) million years ago (MYA); Pavlova *et al.*, 2013), by using multiple genes we improve the precision of the estimate. The optimal partitioning scheme and substitution models were identified using PartitionFinder (Lanfear *et al.*, 2012; Table S3). Linked trees, linked clock models, and unlinked substitution models were used. We performed four replicates with 8 × 10^7^ generations sampled every 2000 steps after 10% of burn-in. The four independent runs were combined and convergence checked in Tracer v1.6.0 (Rambaut *et al.*, 2014). Mitolineage divergence time was calibrated assuming neutral evolution rates for mitochondrial genes of the Hawaiian honeycreeper (Lerner *et al.*, 2011).

## Results

### Nuclear DNA genetic structure

The STRUCTURE models with two and three clusters (*K* = 2, *K* = 3) reached convergence for all chains (Figs. S1-S2). These supported two main clusters (northerly and southerly) with few individuals displaying intermediate assignment scores (red and blue in Fig. 1B and Fig. 2A; Table S1). The higher likelihood model *K* = 3 (Fig. S3) further sub-divided the southern cluster into inland and coastal clusters, in which all inland individuals (Q > 0.8) belong to mito-A mitolineage, and coastal individuals to mito-B mitolineage (shades of blue in Fig. 1B and Fig. 2A). *K* = 4 and K = 5 analyses did not show any additional geographically meaningful structure (not shown). DAPC showed that 48% of genetic variation is explained by the major north-south structure (PC1 on Fig. 2C), and 12% is explained by the minor southerly inland-coast structure (PC2 on Fig. 2C).

### Population divergence: isolation-with-migration model

Convergence of parameter estimates was established by inspecting IMa2 trend plots, and independence of estimates was confirmed by low parameter autocorrelation between steps (mean = 0.013). Posterior parameter distributions reached zero for all parameters, except for migration from the southern to the northern ancestral population (Fig. S4). At least 11,000 genealogies were recorded for each of the eight replicate runs.

IMa2 (Fig. 3A; Table S4) placed divergence between northern and southern populations (*t*1) in the late Pliocene or early Pleistocene (high point 2,400,000; 95% highest posterior density [HPD] 1,900,000 – 3,000,000 years ago), and the split between southern inland and southern coastal populations (*t*0) in the late Pleistocene (64,000; 22,000-118,000 years ago). Compared to the ancestral effective population size (*Ne_ANC_*), sizes of southern (*Ne_S_*) and northern (*Ne_N1_*) populations grew after *t*1, but declined dramatically after *t*0 in all three descendant populations: southern coastal (*Ne_SC_*), southern inland (*Ne_SI_*) and northern (*Ne_N2_*) (Fig. 3A; Table S4). Despite the possibility that gene flow might be under-estimated, we estimated non-zero nuclear gene flow between all ancestral and all current genetic clusters (Fig. 3A; Table S4). For the *t*1-to-*t*0 time period, forward-in-time gene flow from south to north (m2) was higher than that from north to south (m1). For *t*0-to-present, gene flow from southern to northern populations (m4, m8) was higher than that from northern to two southern ones (m3, m7), and gene flow from coast to inland (m6) was higher than that from inland to coast (m5). These nuclear gene flow estimates suggest that neutral gene migration was primarily in the south-to-north direction.

**Figure 3.**
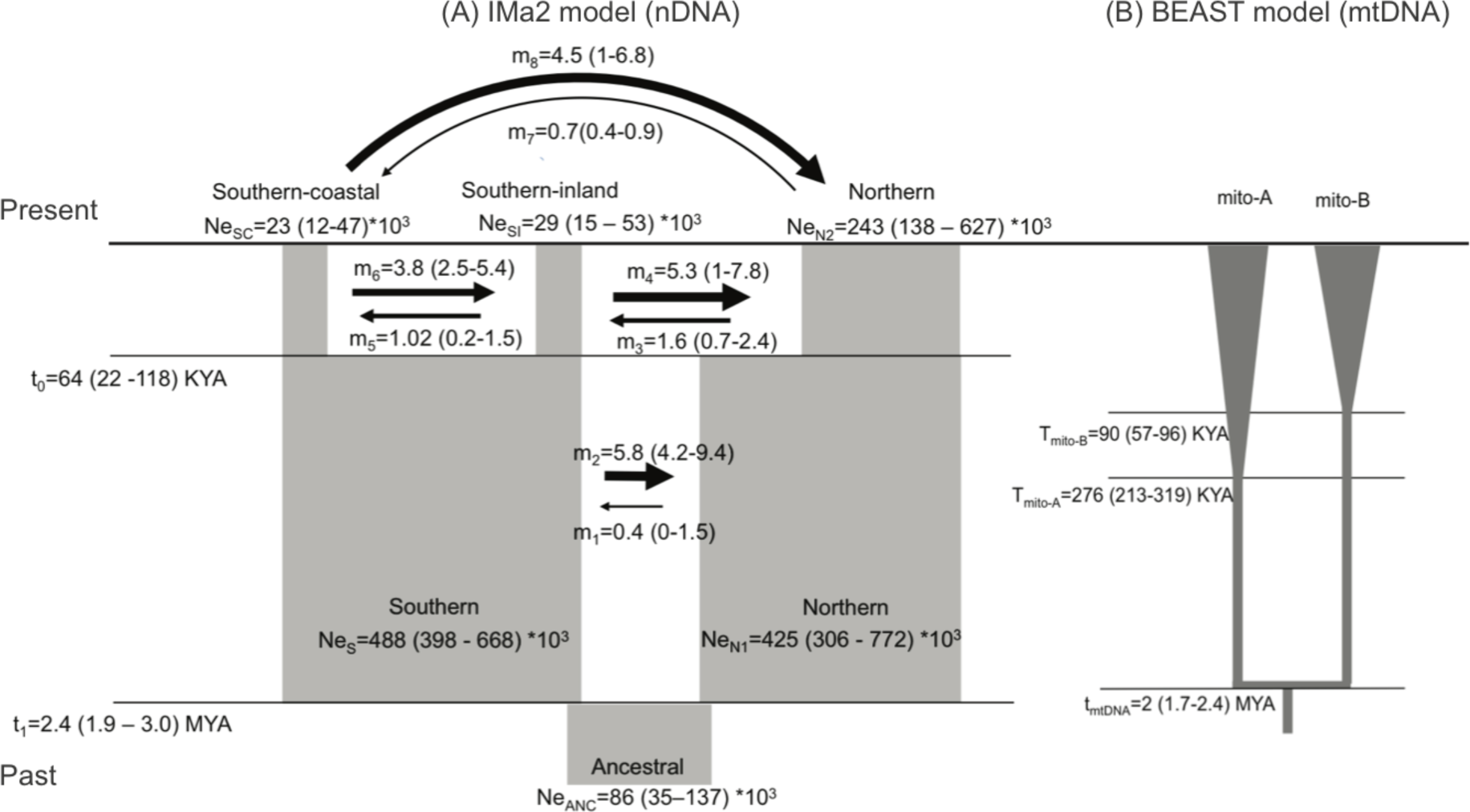
Estimates (high point (95%HPDs)) of coalescent analyses in IMa2 (400 nDNA loci) and BEAST (10 mtDNA genes). (A) IMa2 model. Divergence times: (*t*1) northern vs southern ancestral populations; (*t*0) southern coastal vs. southern inland populations. Effective population size (*Ne*): (*Ne_ANC_*) ancestral root population; (*Ne_N1_*) ancestral northern population; (*Ne_S_*) ancestral southern population; (*Ne_N2_*) northern population; (*Ne_SC_*) southern coastal population; (*Ne_SI_*) southern inland population. Gene flow: (*m*1) ancestral northern to ancestral southern; (*m*2) ancestral southern to ancestral northern; (*m*3) northern to southern inland; (*m*4) southern inland to northern; (*m*5) southern inland to southern coastal; (*m*6) southern coastal to southern inland; (*m*7) northern to southern coastal; (*m*8) southern coastal to northern. (B) BEAST mitolineage model. Divergence times (tmtDNA) between mitolineages mito-A and mito-B. Time to the most recent common ancestor (TMRCA) for (T_mito-A_) mitolineage mito-A and (T_mito-B_) mitolineage mito-B indicate probable times of mitochondrial selective sweeps.

### Mitochondrial lineage divergence

Convergence for the combined BEAST run of the 10 protein-coding mitochondrial gene dataset was confirmed with trend plots and high effective sample sizes (>2000) for all parameters. Mitolineages mito-A and mito-B were estimated to have diverged in the late Pliocene or early Pleistocene (2,000,000; 1,700,000 - 2,400,000 years ago; Fig. 3B; Table S4; Fig. S5). Given that these dates overlap with the 95%HPD of the time of population divergence between northern and southern populations, our result is consistent with the prediction that mitochondrial and nuclear divergence coincided temporally. The time to the most recent common ancestor (TMRCA) for mito-A was placed in the mid Pleistocene (276,000 – HPD: 213,000 – 319,000 years ago) and TMRCA for mito-B in the late Pleistocene (90,000 – HDP: 56,000 – 96,000 years ago). These times are recent relative to the divergence time between mitolineages (Fig. 3B; Fig. S5), presumably because the inland and coastal mitochondrial selective sweeps occurred at this times (Thomson *et al.*, 2000; Rambaut *et al.*, 2008).

## Discussion

Coalescent analyses provided evidence of temporally concordant nDNA and mtDNA divergence. This strongly supports our hypothesis that EYR mitonuclear discordance was caused by population divergence followed by two independent events of mitochondrial introgression. Mitochondrial lineage divergence and introgression generated a deep inland-coastal mitochondrial split in each of the two divergent nuclear genetic backgrounds (north and south; Fig. 1C). This process could have had implications for mitochondrial-nuclear (mitonuclear) associations because divergent inland-coastal mitochondrial types would need to maintain suitable mitonuclear interactions for metabolic functioning under local environmental variation (Burton *et al.*, 2013; Hill, 2015). We suggest that adaptive mitochondrial introgression occurred during a period of transition from relatively warm, stable climates with high summer precipitation, to more variable and winter-dominated rainfall climates, and aridification of inland Australia (Hocknull *et al.*, 2007; Byrne *et al.*, 2008; Sniderman *et al.*, 2009; Byrne *et al.*, 2011). Overall, our data suggest that a first axis of incipient speciation was formed during north-south divergence (reflected in plumage colour subspecies and major nDNA structure), and was supplemented by a second perpendicular axis of incipient speciation during two events of mitochondrial introgression generating inland-coast mitochondrial divergence-with-gene-flow (consistent with two deeply divergent mitolineages).

### Eastern Yellow Robin evolutionary history

The geographic pattern of nDNA, colour variation and mtDNA in EYR indicate that the concerted early-Pleistocene divergence of mtDNA and nDNA occurred in north-south direction. Present-day distributions of nDNA and plumage colour are arranged in a north-south direction (Fig. 1B; Morales et al. submitted). Mitolineage mito-A is the only one currently occurring in the northern part of the species range and displays late-Pleistocene intra-lineage vicariance between the northernmost part of the range and the rest of the mitolineage (Fig. 1A; Fig. S7). Large-scale Pleistocene climatic shifts are likely drivers of the initial north-south divergence (Byrne *et al*., 2008; 2011). Two vicariant/environmental barriers located near the zone of intergradation between northern and southern clusters could have facilitated north-south EYR divergence: the Hunter Valley Barrier and the Southern Transition Zone (Fig. 1B). However, given that contact zones can move (Taylor *et al.*, 2014), the Bassian Volcanic Barrier is another possible candidate (Fig. 1B). These three barriers have been implicated in subspeciation in several other bird species and a variety of other closed-forest taxa including invertebrates, lizards, frogs, mammals and plants (Ford, 1987; Schodde & Mason, 1999; Schodde, 2006; Bryant & Krosch, 2016). Given the clear pattern of north-south historical divergence, to explain the current mitonuclear discordance in EYR we have to invoke two instances of long-range introgression of mitolineages that became fixed without any apparent signs of long-range nuclear introgression. According to this model we can infer the direction and time of introgression by the geographic positions of mitolineages and the estimated time of mitochondrial sweeps (Fig. 1C; Fig. 2B): northern mito-A is inferred to have introgressed southwards along the inland side of the Great Dividing Range in the mid Pleistocene, while southern mito-B introgressed northwards along the coast in the late Pleistocene. Thereafter, in the southern cluster only, nDNA was sorted into coastal and inland populations, in concordance with the mitochondrial split (Fig. 1; Fig. 2A).

### Mitochondrial DNA introgression was likely adaptive

Showing conclusive evidence of fitness effects of mtDNA introgression in wild populations is challenging, but rejecting scenarios of neutral introgression and showing strong genetic evidence of non-neutral evolution are first steps towards demonstrating adaptive nature of mtDNA introgression (e.g. Doiron *et al.*, 2002; Ballard & Melvin, 2010; Boratyński *et al.*, 2014; Llopart *et al.*, 2014). Neutral introgression scenarios under female-biased dispersal and Haldane’s Rule predict higher nDNA than mtDNA introgression and the direction on introgression to be uniform across the entire contact zone (Currat *et al.*, 2008; Excoffier *et al.*, 2009; Rheindt & Edwards, 2011; Toews & Brelsford, 2012). Our results do not support the neutral expectations: (1) massive mitochondrial introgression events in EYR occurred in opposite directions and without much nDNA introgression, (2) the geographic extent of the mtDNA introgression was extreme, and (3) there is previous evidence of adaptive mitochondrial evolution in this system (Pavlova *et al.*, 2013; Morales *et al.*, 2015; Fig. 1). Strong disparities in effective population sizes cannot explain neutral mtDNA introgression because our coalescent estimates show that ancestral populations had equally large population sizes (Toews & Brelsford, 2012). Instead, large ancestral northern and southern population sizes suggest that adaptive mitochondrial variation relevant for lineage bioenergetics might have accumulated at this stage (Fig. 3A; Table 1; Kimura *et al.*, 1963; Ohta, 2002; Camus *et al.*, 2015). Later, adaptive alleles could have fully replaced alternative mitochondrial alleles in response to large-scale climatic change, such as aridification of the inland region, generating the observed selective sweeps (Byrne *et al.*, 2008; Rieseberg, 2009; Byrne *et al.*, 2011; Rheindt & Edwards, 2011). Here we conclude that adaptive mitochondrial introgression in EYR is likely, but further testing of metabolic consequences of introgression requires data on fitness responses to environmental variation and hybridization (e.g. Pereira *et al.*, 2014; Boratyński *et al.*, 2016).

### Perpendicular axes of incipient speciation in the Eastern Yellow Robin

Strong and complex fitness consequences of mitochondrial DNA variation (Wolff *et al.*, 2016) are likely to be further amplified through mitonuclear interactions (Bar-Yaacov *et al.*, 2012; Deremiens *et al.*, 2015; Boratyński *et al.*, 2016). Mitonuclear co-evolution is essential for critical metabolic and physiological cell functions. Disrupted mitonuclear interactions can form strong, long-lasting barriers to gene flow and promote speciation (Dowling *et al.*, 2008; Gershoni *et al.*, 2009; Burton *et al.*, 2013; Hill, 2015; Hill, 2016). In EYR, one possible explanation for the dramatic reduction of effective population sizes in all three populations at (*t*0) is strong selection against hybrids bearing incompatible mitonuclear combinations, after the onset of mitochondrial introgression (Fig. 1C; Fig. 3A). Evidence from laboratory crosses across a wide range of animal systems shows that mitonuclear incompatibility fitness effects in hybrids include metabolic malfunctioning, low fertility and increased mortality (reviewed in Burton *et al.*, 2013; Levin *et al.*, 2014). Under this incompatibility theory, mitonuclear functioning would be expected to ameliorate over time through co-introgression and further refinement of co-evolved mitonuclear genes (Beck *et al.*, 2015) and be maintained by selection against mitonuclear mismatches (Burton *et al.*, 2013). Testing these predictions will require analysis of genome-wide variation including a thorough inspection of mitonuclear genes (Bar-Yaacov *et al.*, 2015). Additionally, mitonuclear co-evolution in coastal and inland lineages might be maintained by female choosiness of males with compatible mitonuclear genes (i.e. the mitonuclear sexual selection hypothesis; Hill & Johnson, 2013). According to this theory, males can advertise their mitonuclear type with signals linked to mitochondrial metabolism (such as plumage colour or song). Despite that both northern and southern nDNA clusters contain divergent EYR mitolineages, we identified nuclear genetic substructure concordant with mitolineage membership only in the southern cluster (Fig. 1B, Fig. 2). A southward mitochondrial introgression of mito-A could have occurred earlier than northward introgression of mito-B according to TMRCA estimates, aligning with the time estimate for mitochondrial selective sweeps (Fig.1C; Fig. 3B). Thus, the southern minor nDNA structure might reflect stronger selection against gene flow in the south than in the north due to longer evolution of reproductive barriers. Behavioural experiments are required to test whether EYR mate assortatively according to mitolineage (Hill & Johnson, 2013).

Despite the large amount of data amassed for EYR, it remains unclear whether either or both of the perpendicular axes of incipient speciation will lead to full speciation. Our data show the importance of understanding phenotypic and mitonuclear diversity patterns before jumping to taxonomic conclusions. Speciation is a complex process that operates along a continuum, and under the biological species concept any trajectory towards full speciation in EYR seems to been halted for now (Seehausen *et al.*, 2014; Shaw & Mullen, 2014). The real challenge for taxonomy then may be that of upon which criteria one can meaningfully, or indeed whether one should, diagnose any intraspecific populations e.g., on plumage and nDNA, or on mtDNA. On the other hand, biological and conservation implications of EYR intraspecific variation are clearer: evidence for natural selection operating in different directions and evidence for restricted gene flow on historical timescales indicate the existence of genetically and ecologically interchangeable EYR populations (Crandall *et al.*, 2000).

### Speciation drivers in Eastern Australia

Detailed reconstruction of Quaternary climates in Australia is limited by a paucity of suitable fossil sites, a relatively narrow range of modern climate space for calibration, and large regional variability (Hocknull *et al.*, 2007; Sniderman *et al.*, 2009; Porch, 2010; Sniderman, 2011; Saltré *et al.*, 2016). However, there is consensus on key paleoclimatic phenomena that could have driven speciation in EYR in two perpendicular axes. The most important phenomena are: (i) a major transition in the early Pleistocene from relatively warm, stable climates with high summer precipitation, to later more variable ones characterized by winter-dominated rainfall and, (ii) major differences between northern and southern Australia in the severity and timing of increased summer aridity, and (iii) more severe aridity inland than on the coast (Hocknull *et al.*, 2007; Byrne *et al.*, 2008; Sniderman *et al.*, 2009; Byrne *et al.*, 2011).

Multiple lines of evidence indicate that warm and moist early-Pleistocene climates in Australia persisted until transition to modern winter-dominated rainfall (Sniderman *et al*., 2009). These conditions are likely to have applied broadly across south-eastern Australia, and would have been in place during our proposed phase of north-south divergence of EYR. The adaptive introgression that we propose occurred would have done so during a transition from the mild early-Pleistocene climate to more variable climates with winter-dominated rainfall and summer aridity. This period of climate change involved considerable temporal and spatial complexity, such as oscillation between wetter and drier vegetation types and multiple periods of rapid fall in humidity and temperature (Hocknull *et al.*, 2007; Sniderman *et al.*, 2009; Sniderman, 2011; Saltré *et al.*, 2016). However, it is clear that this shift occurred later in northern Australia than in the south: a north-eastern tropical rainforest fauna 500-280 KYA was replaced by a xeric fauna 205-170 KYA, while the south experienced increasing aridity from as early as 600 KYA (Hocknull *et al*., 2007). In addition, due to the rain-shadow effect of the Great Dividing Range, aridity is more severe inland than on the coast. This pattern also likely developed earlier in southern than northern Australia (Hocknull *et al*., 2007).

We predict that many co-distributed species may have been impacted similarly by these widespread paleoclimatic events. If adaptive mitochondrial introgression is common in the Australian avifauna (e.g. Kearns *et al.*, 2014; Shipham *et al.*, 2015), then that, coupled with strong shifts in climate-related selective forces, could provide a general explanation for the very high rate of mitochondrial paraphyly observed in Australian birds (Joseph & Omland, 2009). Accordingly, strong shifts in environmental gradients associated with paleoclimatic cycling could also be a common general mechanism for mitonuclear co-evolution as a driving force in generating genomic conflict and mitonuclear speciation (Hill, 2015). Whether similar phenomena have occurred in other taxa, and on other continents with suitable conditions, demands further investigation of evolutionary impacts and biodiversity implications of Quaternary climate change.

## Acknowledgements

Funding was provided by Australian Research Council Linkage Grant (LP0776322) and the Holsworth Wildlife Research Endowment (2012001942). HM was funded by a Monash University with a Graduate Scholarship (MGS), a Faculty of Science Dean’s International Postgraduate Research Scholarship, and a Postgraduate Publication Award (PPA) and by the Department of Public Education (SEP) of the Mexican Government. Bioinformatic analyses were undertaken using the Monash Sun Grid high-performance computer facility. We are grateful to Philip Chan for technical support. Field samples were collected afresh for this study under scientific research permits issued by the Victorian Department of Environment and Primary Industries (numbers 10007165, 10005919 and 10005514) and New South Wales Office of Environment and Heritage (SL100886). We thank Nevil Amos and Richard Major for coordinating the fieldwork and Holly Sitters and Christine Connelly for providing genetic samples from South and East Victoria. We thank Anders Gonçalves da Silva and Biao Wang for inputs regarding data analysis, and Kaspar Delhey for valuable discussions that help to improve the manuscript. We thank Scott Edwards, Mike Webster, Lynna Kvistad and Stephanie Falk for valuable comments on the manuscript. We are grateful to three anonymous reviewers and editor Matthew Miller from Axios Review for helpful comments on the first draft of this manuscript. The authors are also grateful to Alan Lemmon for coordinating the hybrid capture sequencing project.

## Appendix

Targeted capture sequencing methods

STRUCTURE analysis methods

Table S1 Samples screened for nuclear DNA (nDNA) and mitochondrial ND2 variation

Table S2 Samples screened for mitochondrial genome

Table S3 Partitions for the BEAST analysis and substitution models according to PartitionFinder

Table S4 Parameter estimates for coalescent analyses in BEAST and IMa2

Table S5 Speciation hypotheses in the Eastern Yellow Robin

Figure S1 Convergence plots for STRUCTURE K = 2

Figure S2 Convergence plots for STRUCTURE K = 3

Figure S3 Delta likelihood for STRUCTURE analysis with clusters *K* = 1-5

Figure S4 Posterior distributions for IMa2 model

Figure S5 Phylogenetic tree reconstructed by BEAST from 10 mitochondrial genes

### Data access in doi:10.6084/m9.figshare.3581004

Sample Information

Hybrid capture probes design

BLAST output file

Fasta alignment files of nDNA loci

SNP data

Alignment of 10 mtDNA genes in nexus format

